# Ectopic Expression of Virus Innexins in *Heliothis virescens* Disrupts Hemocyte-Mediated Encapsulation and Host Viability

**DOI:** 10.1101/788661

**Authors:** Peng Zhang, Matthew W Turnbull

## Abstract

1.

Polydnaviruses are dsDNA viruses associated with endoparasitoid wasps. Delivery of the virus during parasitization of a caterpillar and subsequent virus gene expression is required for production of an amenable environment for parasitoid offspring development. Consequently, understanding of Polydnavirus gene function provides insight into mechanisms of host susceptibility and parasitoid wasp host range. Polydnavirus genes predominantly are arranged in multimember gene families, one of which is the *vinnexins*, which are virus homologues of insect gap junction genes, the *innexins*. Previous studies of *Campoletis sonorensis* Ichnovirus Vinnexins using various heterologous systems have suggested the four encoded members may provide different functionality in the infected caterpillar host. Here, we expressed two of the members, *vnxG* and *vnxQ2*, using recombinant baculoviruses in susceptible host, the caterpillar *Heliothis virescens*. Following intrahemocoelic injections, we observed >90% of hemocytes (blood cells) were infected, producing recombinant protein. Larvae infected with a *vinnexin*-recombinant baculovirus exhibited significantly reduced molting rates relative to larvae infected with a control recombinant baculovirus and mock infected larvae. Similarly, larvae infected with *vinnexin*-recombinant baculoviruses were less likely to molt relative to controls, and showed reduced ability to encapsulate chromatography beads in an immune assay. In most assays, the VnxG protein was associated with more severe pathology than VnxQ2. These results, in light of previous findings, support that Polydnavirus Vinnexin gene family members may provide complementary, rather than redundant, effects. This in turn indicates a need to test gene family member functionality across infected hosts for effects to determine member contribution to host range.

**Importance:** Polydnaviruses are obligate mutualistic associates of highly speciose wasp taxa that parasitize caterpillars. Expression of Polydnavirus-encoded genes in hosts parasitized by wasps is necessary for successful parasitization, and an unusual genome structure including multiple-membered gene families is hypothesized to contribute to host manipulation. We have tested this hypothesis by *in vivo* expression of two members of a family of Polydnavirus homologues of Innexins, or insect gap junction proteins. Previous findings demonstrated that the two Vinnexins induce different physiological alterations in heterologous systems. Here, in host caterpillars, we observed differential alteration by the two proteins of host immune cell (hemocyte) bioelectrical physiology and the immune response of encapsulation. Not only do our data suggest a linkage between cellular bioelectricity and immunity in insects, but they support that gene family expansion has functional consequences to both Polydnavirus and host wasp success.

## 3. Introduction

Polydnaviruses (PDVs) are a remarkable virus family. These large dsDNA viruses are obligately and mutualistically associated with specific families of parasitoid wasps, a relationship that generates unusual selection pressures on the viruses. The association of these viruses with their wasp hosts is essential to the ecological and evolutionary success of those wasps. PDVs are actually a paraphyletic linage: Bracoviruses (BVs), associated with PDVs are associated with microgastroid complex of the wasp family Braconidae, while Ichnoviruses (IVs) are associated with Campopleginae and Banchinae wasp subfamilies of the wasp family Ichneumonidae. Despite their evolutionary independence, the different lineages of PDVs exhibit similar genome organization and are integrated as a provirus into the genome of the host wasp. The genome of members of both lineages is comprised by large dsDNA across multiple segments, of which a majority of the sequence is non-coding, while coding sequences cluster into numerous multimember gene families (1, 2). The PDV genome is transmitted vertically as a provirus in the wasp genome, while horizontal transmission occurs across species boundary: PDV replication and encapsidation precedes oviposition, during which virus is introduced along with egg into the hemocoel of a parasitized insect (typically caterpillar) host. Infection followed by expression of virus genes results in numerous host pathophysiologies, including disruptions of the immune, energy homeostatic and endocrine systems, which provide a beneficial environment to the developing wasp (2–4). There is no evidence of PDV replication in the infected secondary host during this period (5).

As virus replication does not occur in the caterpillar host, it generally has been held that PDV genes expressed in that host play a role in generating pathophysiology (3, 6, 7). Therefore, expansion of PDV genes into gene families has been viewed as evidence of selection for manipulation of additional hosts and/or tissues and processes within a host (8, 9). Assigning function to PDV genes then provides insight into the mechanisms by which PDVs manipulate the insect host of the PDV, i.e., the mechanisms by which the associated parasitoid manipulates the host. However, due to the complexities of the association between wasp and virus, it can be difficult to assess the roles of individual genes and/or gene families. Thus, work in both BVs and IVs has sought to elucidate PDV gene function, typically testing sufficiency using *in vitro* assays or necessity via RNAi during PDV infection (10–14).

Total IV gene expression is associated with multiple host pathologies, such as alterations of cytoskeleton (15–17) and disruption of cell signaling (10). IV protein functional analyses have mostly focused on the cys-motif family found in all IV genomes to date (18–20), especially its role in suppressing immune system and manipulating host translation (21–23). The vinnexins are another IV multimember gene family (24). These genes are virus homologues of insect gap junction genes (*innexins*) and are highly conserved among IVs (25). The proteins form functional gap junctions (24, 26), and *in vitro* expression of *vinnexins* induces cell membrane depolarization and cytoplasmic alkalization (27). Results to date imply the Vinnexins have protein-specific functions (26–28). As Innexins occur and function in hemocytes including during immune responses (29–32), and gap junction disruption reduces immunocompetence (28), we hypothesize the Vinnexins may interfere with immune responses. The ability of individual Vinnexins to differentially affect hemocytic immunity *in vivo* has not been thoroughly tested.

To develop a better understanding of the effects of Vinnexins on larval lepidopterans, we used recombinant baculoviruses to express two vinnexins of the Polydnavirus *Campoletis sonorensis* Ichnovirus (CsIV), *vnxG* and *vnxQ2*, in host *Heliothis virescens* caterpillars. Our results demonstrate that Vinnexin-expressing caterpillars exhibit stunted development and increased mortality relative to a control recombinant baculovirus. Although vinnexin expression does not alter total hemocyte numbers relative to the control baculovirus, both Vinnexins are associated with hemocyte membrane depolarization. VnxG and VnxQ2 differentially affect immunocompetence of infected caterpillars, as VnxQ2 reduces cellular immunity, while VnxG decreases consistency of immune response. Taken together, our data provide further evidence that the Vinnexin gene family members may have different roles within hosts.

## 4. Results

### *In vivo* recombinant protein expression

We utilized recombinant baculoviruses that were previously characterized (27). These include two experimental recombinant viruses that encode *Campoletis sonorensis* Ichnovirus (CsIV) *vinnexinG* (*vnxG*) and CsIV *vinnexinQ2* (*vnxQ2*), each with C-terminus 6x-His epitope, and a third recombinant virus encoding a fish melanocortin-4-receptor (Mc4r) with an N-terminus FLAG epitope to serve as control for effects of recombinant virus and overexpression of an exogenous membrane protein. Newly molted 4th instar *H. virescens* larvae were anesthetized and injected with 100 plaque-forming unit (pfu) of recombinant viruses encoding *mc4r*, *vnxG* or *vnxQ2*, or the same volume of virus-free media. Hemocyte protein was isolated and analyzed by anti-His western blot, which verified recombinant protein expression in hemocyte samples (**Fig. 1A**). We also examined protein expression in hemocytes using epifluorescence immunomicroscopy. Anti-His (**Fig. 1B**) and anti-FLAG immunomicroscopy results (**Fig. 1C**) corresponded to patterns observed with expression of VnxG-His, VnxQ2-His, and FLAG-Mc4r in infected Sf9 cells (27). We simultaneously probed with an antibody against the AcMNPV envelope protein GP64 to determine rate of AcMNPV infection, and typically observed rates in excess of 90%.

**Fig. 1.**
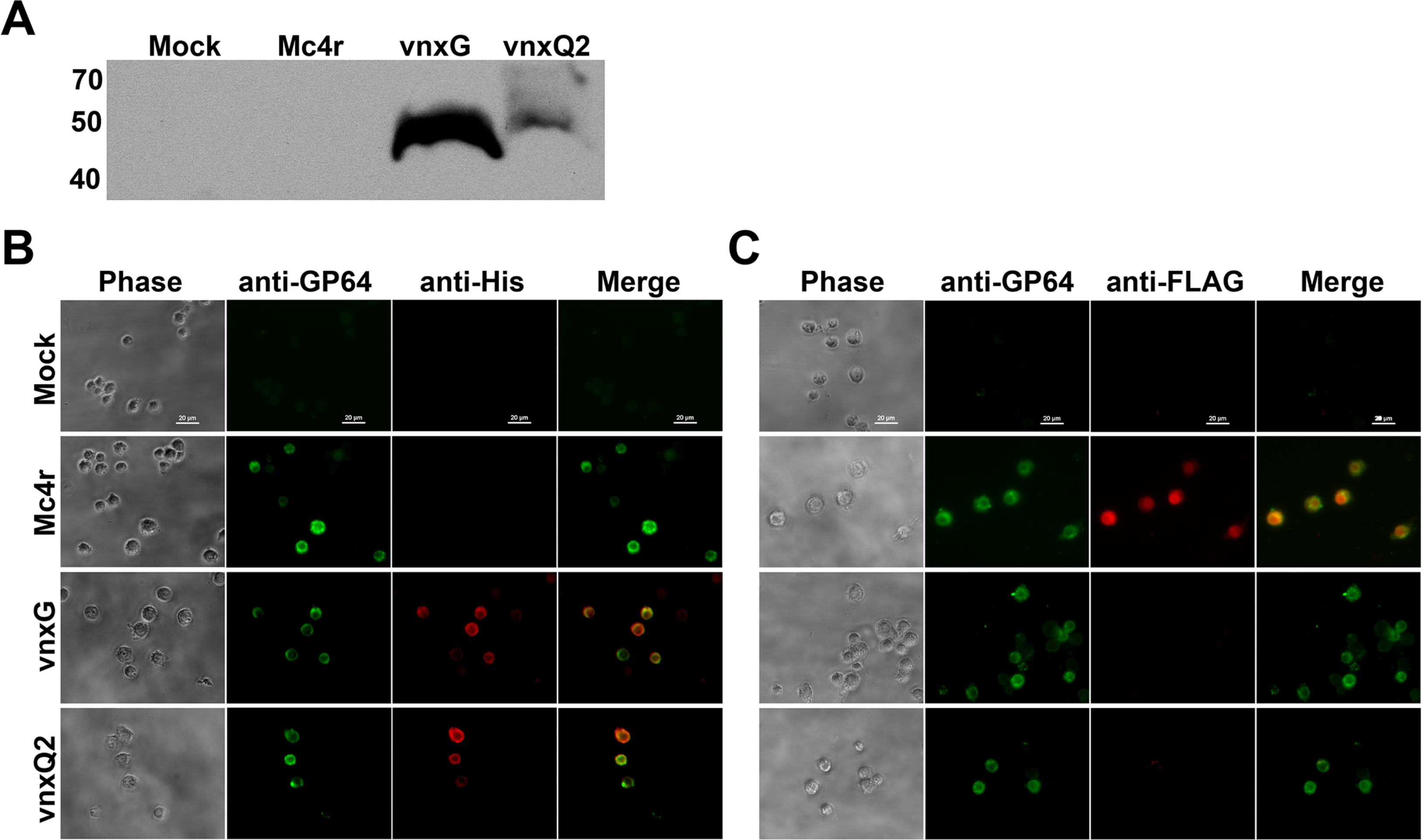
*Vinnexins* are expressed by recombinant baculoviruses in *H. virescens* larval hemocytes. Newly molted 4th instar caterpillars injected with 100 pfu recombinant baculoviruses or media and were bled for hemocyte sample at 3 dpi. (A) Expression of *vinnexin* was verified with western blot. Mock: Mock Treated, Mc4r: AcMNPV-FLAG-*m*c4r, VnxG: AcMNPV-vnxG-His, VnxQ2: AcMNPV-vnxQ2-His. (B) Anti-His and Anti-GP64 immunomicroscopy was performed to verify Vinnexin expression and AcMNPV infection, respectively. (C) Anti-His and Anti-FLAG immunomicroscopy was performed to verify FLAG-Mc4r expression and AcMNPV infection.

### Infection with vinnexin recombinant baculoviruses increases caterpillar mortality

*H. virescens* is a permissive host to wild type AcMNPV, the baculovirus we used for recombinant virus generation. We tested whether recombinant vinnexin expression altered mortality in the presence of an AcMNPV infection. Media or media plus 100 pfu budded control or experimental recombinant AcMNPV was injected intrahemocoelically, and larval mortality was assessed daily for 7d. Approximately 10% of mock infected (media) larvae died during the period, while the control Mc4r virus treatment resulted in ~40% mortality and both Vinnexin recombinants resulted in >85% mortality (**Fig. 2**). Bonferroni correction was conducted ahead of multiple comparison (α’=0.0083). AcMNPV-Mc4r induced significantly higher mortality relative to Mock control (p<0.008). Infection with either *vinnexin* recombinant virus resulted in significantly greater mortality than the Mc4r recombinant virus (p<0.008), although there was no difference between the VnxG and VnxQ2 infection results (p=0.43).

**Fig. 2.**
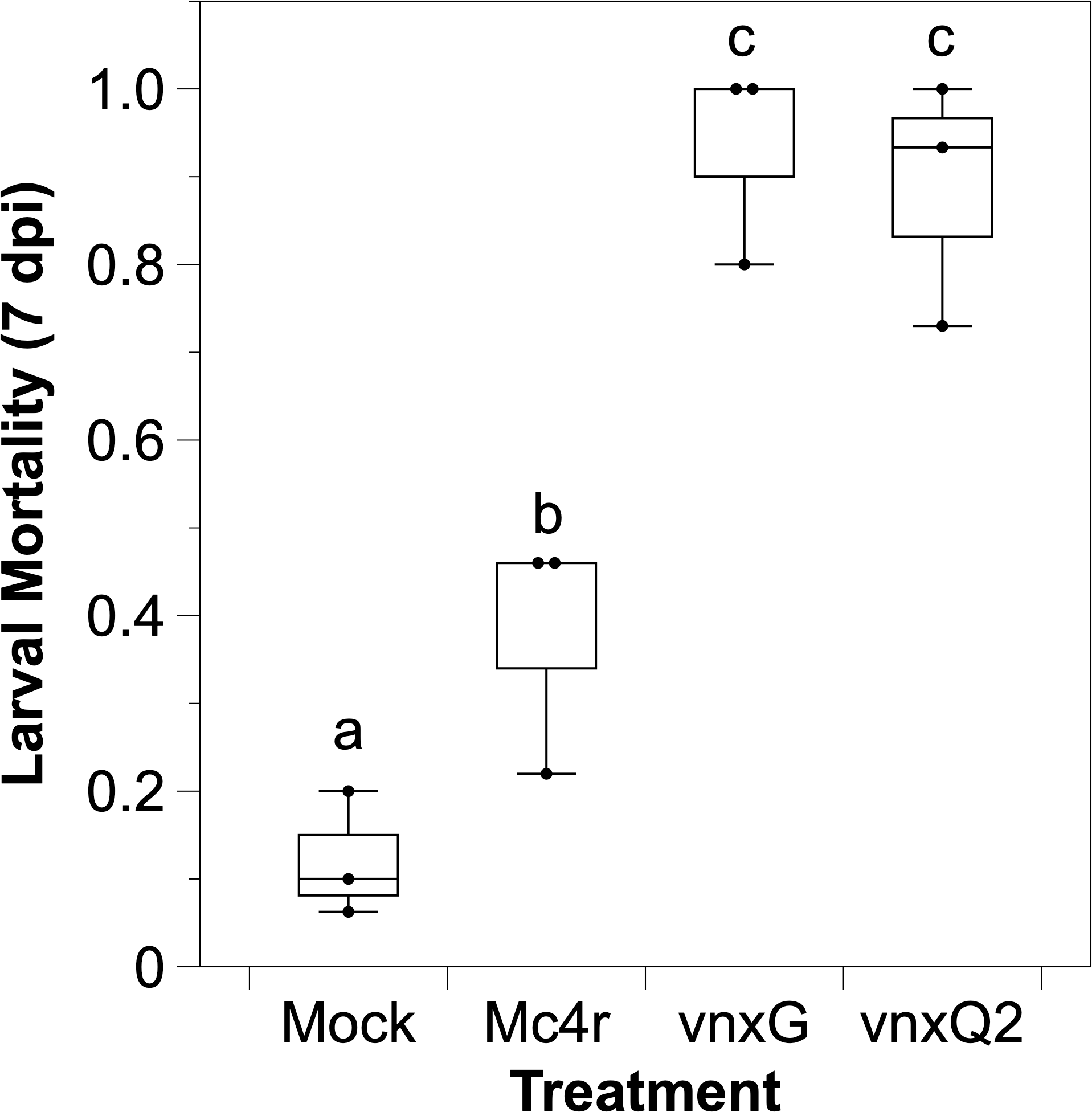
*Vinnexin* expression does not affect total hemocyte count of *H. virescens* larvae. Hemocytes were collected at 3 dpi from 3 caterpillars and were counted with hemocytometer. The unit of cell count is ten million per milliliter. Bars with different letter indicate significant difference at *p* <0.05. Error bars show standard deviation.

### Infection with vinnexin recombinant baculoviruses slows development of host caterpillar

Next, the consequence of Vinnexin expression on developmental rate was examined. A significant difference in proportion of larvae molting from 4_th_ to 5_th_ instar was observed (**Fig. 3**, χ^2^ (3, N=4) = 58.21, p<0.01). Larvae injected with 100 pfu of control AcMNPV-Mc4r did not significantly differ from larvae injected with media in the percentage molting from 4_th_ to 5_th_ instar by 3 dpi (p=0.24). However, both *vinnexin* recombinants differed significantly from both media and AcMNPV-mc4r treatments, with significantly more of the vnxG-infected and vnxQ2-infected larvae failing to molt to 5_th_ instar. Furthermore, there was a significant difference between vnxG-infected and vnxQ2-infected (p=0.02), with significantly fewer AcMNPV-vnxG-injected larvae molting.

**Fig. 3.**
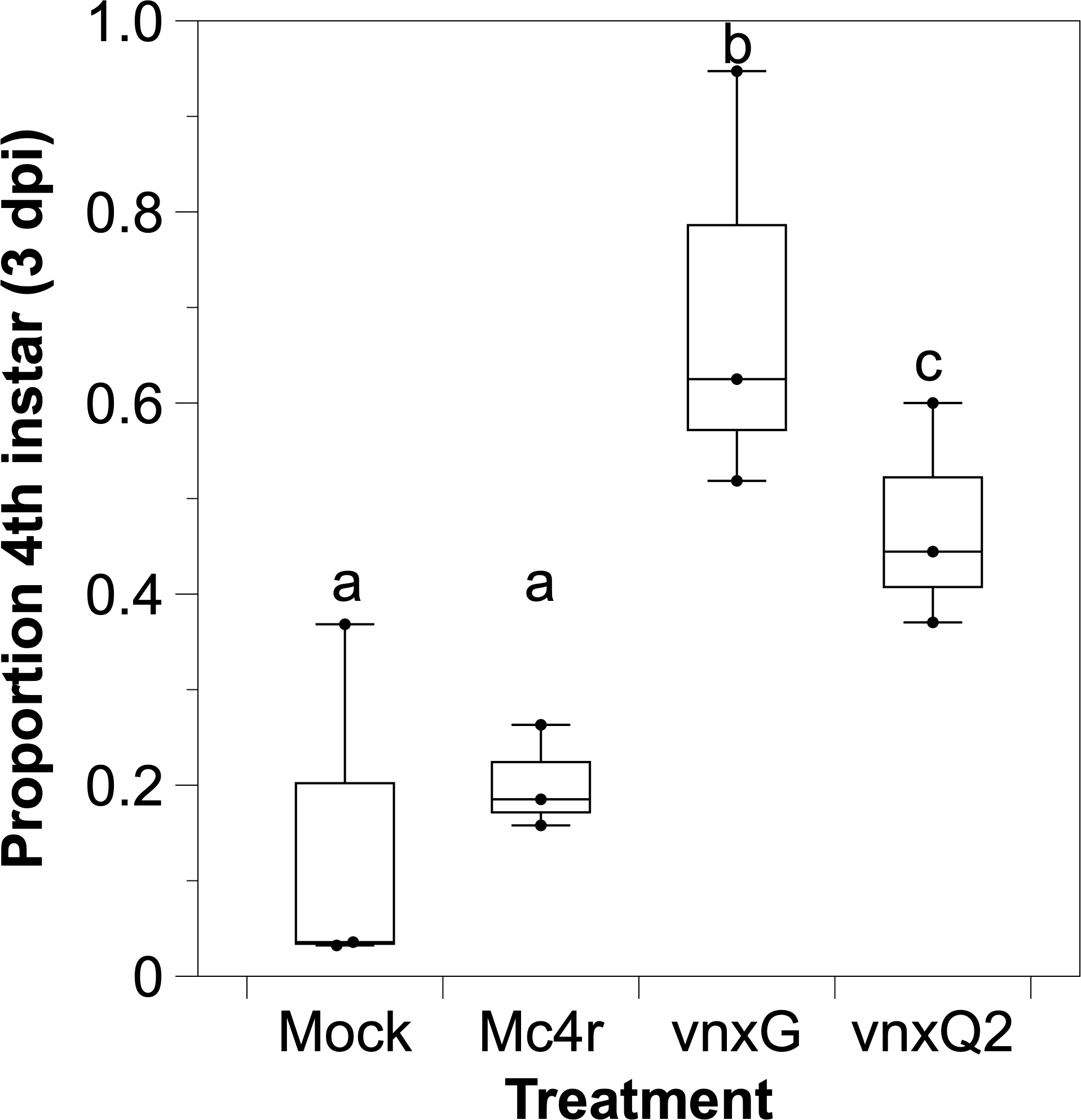
*Vinnexin* expression leads to a higher mortality during NPV infection. Young 4th instar larvae were injected with 100 pfu recombinant viruses or media and maintained at room temperature. Infected caterpillars were inspected daily, and mortality was recorded for one week. The caterpillar mortality for each treatment was compared. Superscript letters indicate significance after Bonferroni correction (α’=0.0083). Error bars show standard deviation.

### Vinnexin expression has no effect on Total Hemocyte Counts (THC)

Previously, we demonstrated that Sf9 cells infected with AcMNPV-vnxG and AcMNPV-vnxQ2 exhibited a significant reduction in cell number relative to Mock-infected and AcMNPV-mc4r-infected cells (27). As a high percentage of hemocytes of larvae injected with the three viruses were GP64 positive, indicating hemocytes were infected with the viruses, we tested whether infection likewise altered hemocyte number *in vivo*. Although there was a trend to reduced THC in virus infected larvae relative to mock infected, the reduction was not significant (ANOVA, F (3,8) =0.21, p =0.89) (**Fig. 4**). There was no significant difference between the THC of Vinnexin-infected and Mc4r-infected larvae (p >0.90) or the VnxG-infected and VnxQ2-infected larvae (p >0.99).

**Fig. 4.**
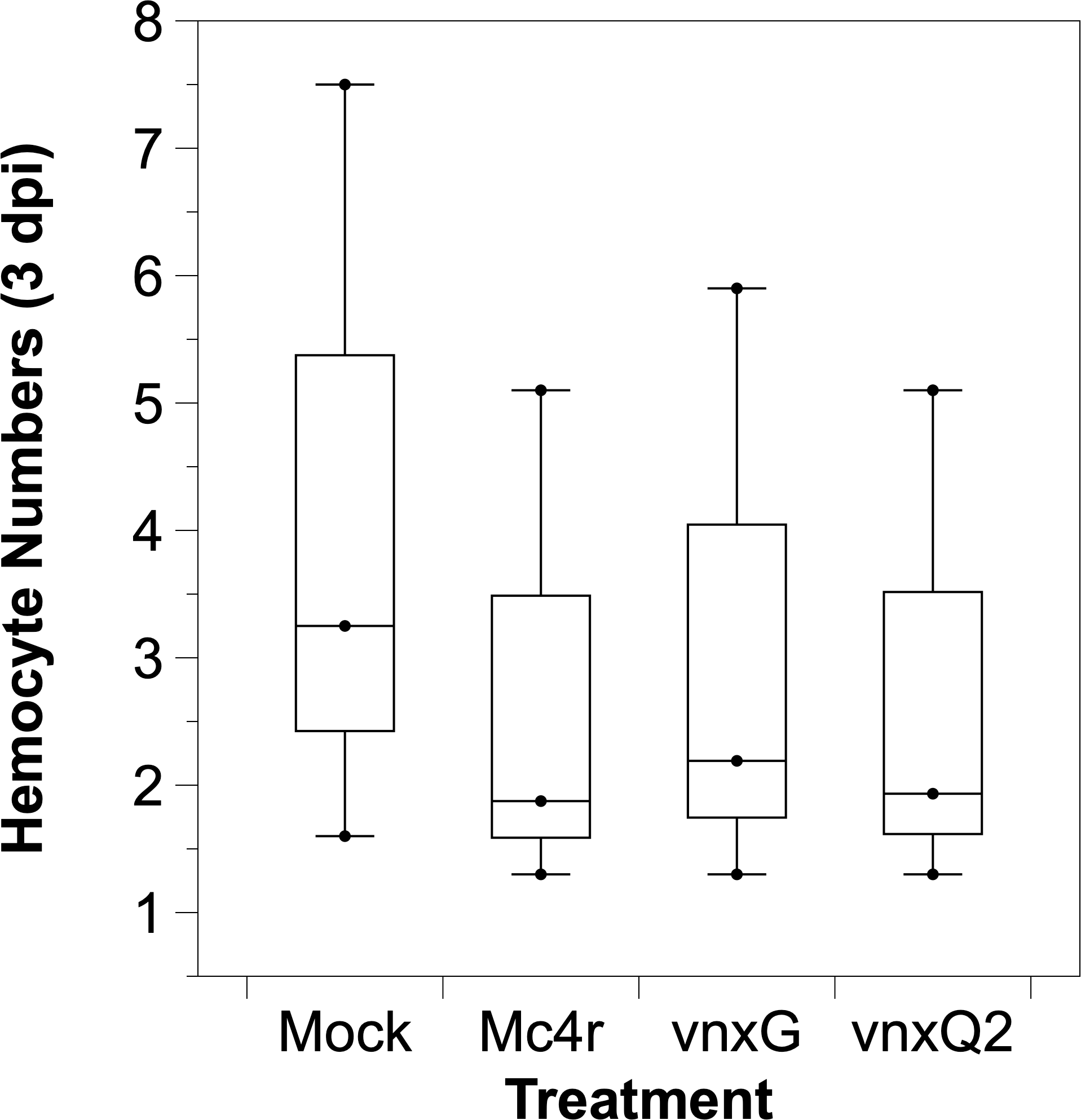
*Vinnexin* expression delays caterpillar development. Mock and virus infected caterpillars were maintained and observed for 3 d post-infection. The developmental stage for individual caterpillars was recorded daily as 4th instar or 5th instar. The proportion of 4th instar larvae was calculated and compared. Superscript letters indicate significant difference at *p* <0.05. Error bars show standard deviation.

### Vinnexin expression depolarizes cell membranes

Membranes of Sf9 cells infected with AcMNPV-vnxG or AcMNPV-vnxQ2 were found to be significantly depolarized relative to controls in a previous study (27). To test for this phenomenon *in vivo*, we isolated hemocytes from mock- and recombinant AcMNPV-infected larvae and incubated them with the membrane potential sensitive dye DiBAC4(3). Cells were manually outlined, and area and normalized intensity determined. Cell area differed significantly between treatments (ANOVA, F(3,1116) =36.53, p < 0.01) (**Fig. 5A**). While AcMNPV-mc4r infection did not alter area relative to mock (p = 0.99), hemocytes from both AcMNPV-vnxG (p < 0.01) and AcMNPV-vnxQ2 (p < 0.01) infected larvae were significantly smaller than those of AcMNPV-Mc4r control. Hemocytes isolated from vnxG-infected larvae were significantly smaller than those from the vnxQ2-infected larvae (p < 0.01).

**Fig. 5.**
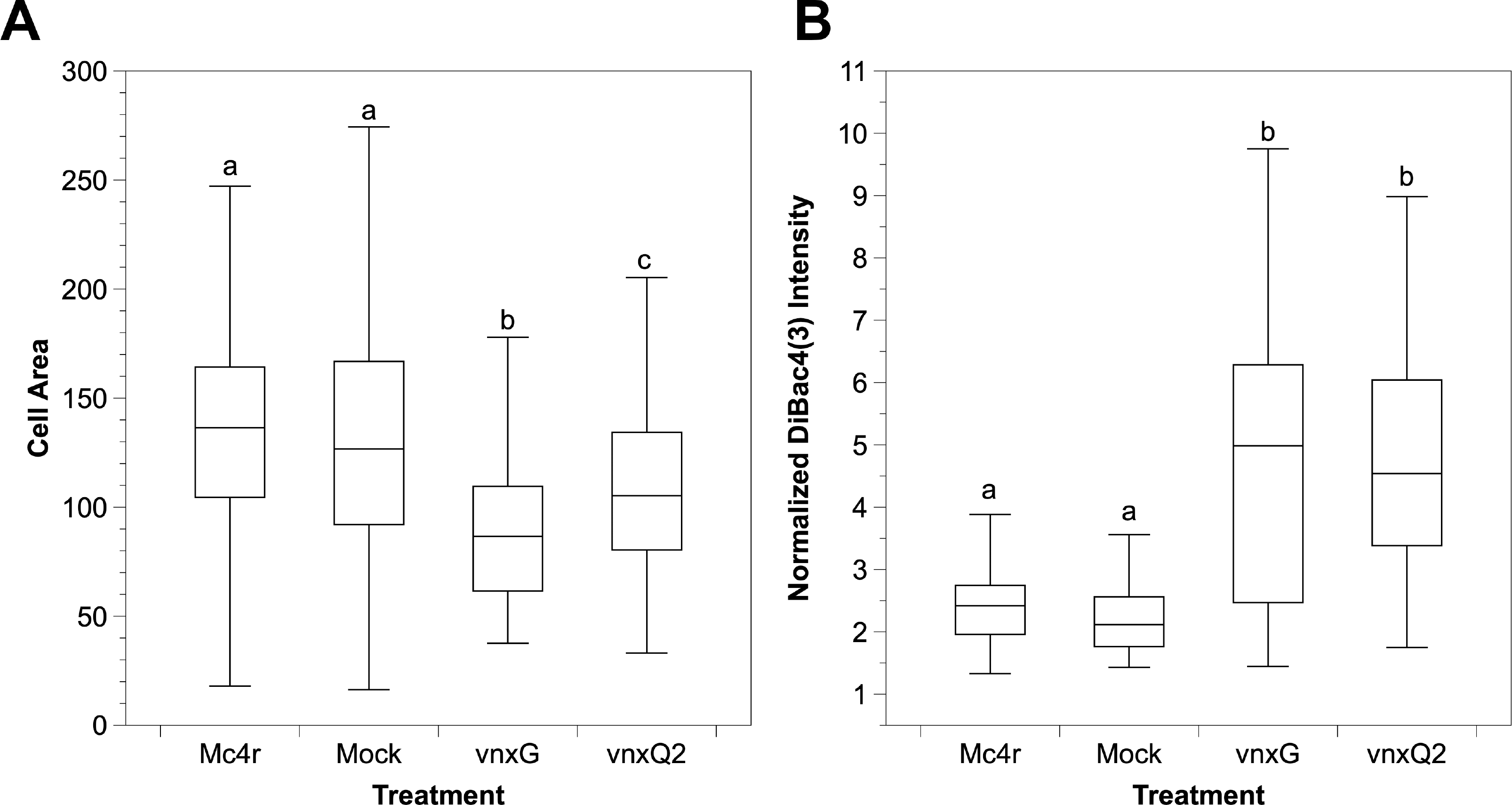
*Vinnexin* expression causes depolarization of hemocytes. Hemocytes of each treatment were incubated with media containing 1μg/ml DiBAC4(3) at 3 dpi. (A) Area of hemocytes. (B) Normalized intensity shows *vinnexin* expression (vnxG or vnxQ2), but not mock treatment or control virus (Mc4r), induces membrane depolarization. Superscript letters indicate significant difference at *p* <0.05.

Normalized DiBAC4(3) intensity (i.e., V_mem_) was significantly affected by recombinant virus infections relative to mock (ANOVA, F(3,1116) = 121.6, p < 0.01) (**Fig. 5B**). AcMNPV-Mc4r infection significantly increased normalized DiBAC4(3) relative to mock (p = 0.01), while AcMNPV-vnxG (p < 0.01) and AcMNPV-vnxQ2 (p < 0.01) significantly increased normalized DiBAC4(3) values in comparison to the AcMNPV-Mc4r control. Hemocytes from AcMNPV-vnxG infected individuals exhibited significantly higher normalized DiBAC4(3) values than those from AcMNPV-vnxQ2 (p < 0.01). Thus, our data indicate that infection with a Vinnexin-expressing AcMNPV induces membrane depolarization in hemocytes, similar to our previous findings in Sf9 cells (27).

### Vinnexin expression disrupts hemocyte encapsulation function

The impact of Vinnexin expression on larval immunocompetence was assessed using an *ex vivo* encapsulation assay. Hemocytes were isolated from mock or rec-AcMNPV infected larvae 3d post-injection and challenged with chromatography beads in an agarose-lined well. Hemocytes from mock- and AcMNPV-Mc4r-injected individuals typically encapsulated beads: >80% of beads were surrounded by several layers of hemocytes, forming a regular shaped capsule (2 on the scale used), while <20% of beads showed irregular or no encapsulation (1 on scale) (**Fig. 6A-B**). Hemocytes from AcMNPV-vnxQ2-injected individuals failed to encapsulate nearly 50% of the presented beads. Intriguingly, hemocytes from AcMNPV-vnxG-injected individuals seldomly formed regular capsules (6%), instead failing to form a capsule or forming a partial one (1 on scale) around >50% of their target beads, and forming larger and more intensive capsules around the remaining 40+% of beads (3 on scale).

**Fig. 6.**
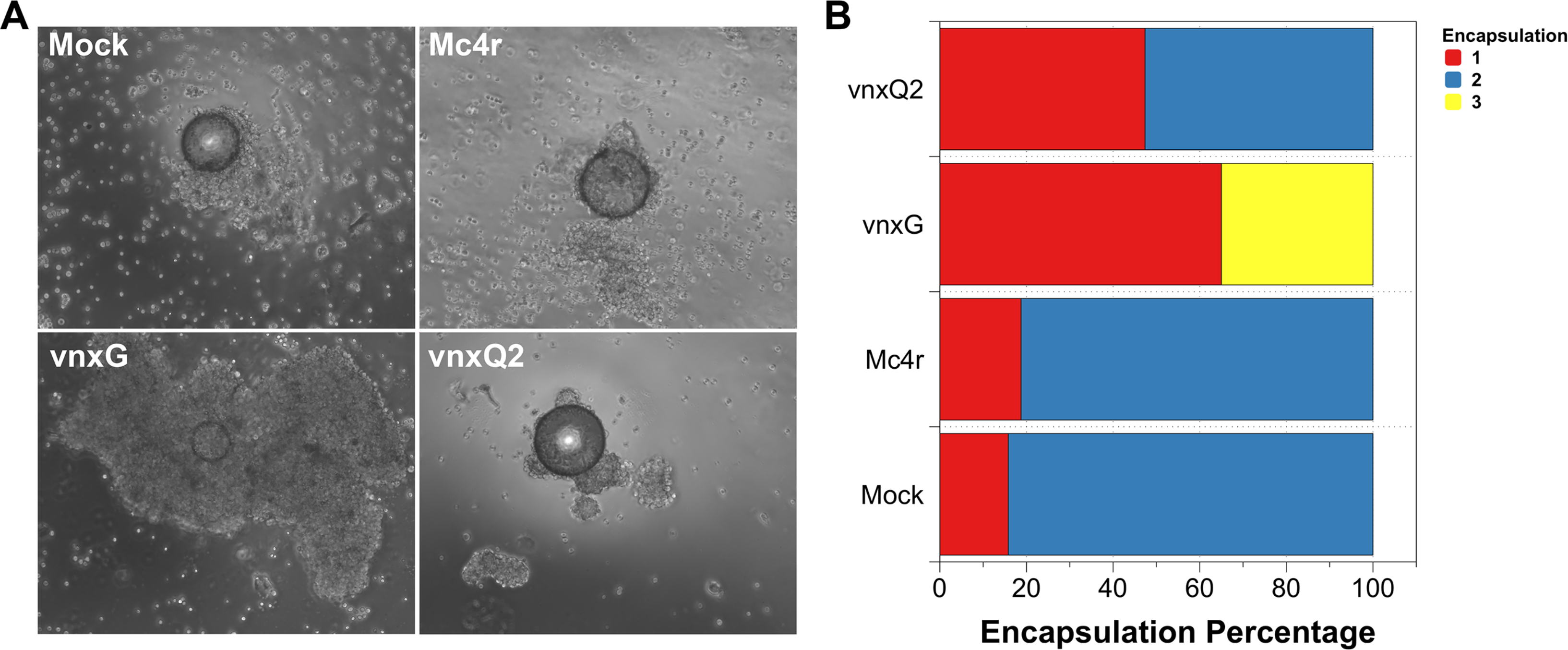
*Vinnexin* expression disturbs hemocyte encapsulation function. *In vitro* encapsulation assay results demonstrate only the expression of VnxG induces atypically large capsule formation, while the expression of VnxQ2 results in reduced encapsulation rate. (A) Representative micrographs of encapsulation results for each treatment. (B) Percentage of encapsulation shown by hemocytes of each treatment according to a 1-3 scale (see text).

For statistical purposes, partial and intensive encapsulations were classified as “abnormal encapsulation” as opposed to “normal encapsulation”. Abnormal encapsulation rate was calculated and compared among the four treatments (χ^2^ = 40.637, df = 3, p < 0.05). Multiple comparison was conducted after Bonferroni correction (α’ = 0.0083). While the two controls did not differ significantly in percentage of beads encapsulated (χ^2^ = 0, p = 1.0), AcMNPV-vnxG differed significantly from both Mock treated (χ^2^ = 30.767, p < 0.0083) and NPV-Mc4r (χ^2^ = 30.767, p < 0.008). Although AcMNPV-VnxQ2 had more partial encapsulation, the proportion was not significantly different from either Mock (χ^2^ = 3.857, p = 0.05) or AcMNPV-Mc4r (χ^2^ = 3.857, p = 0.05).

## 5. Discussion

The relationship of PDVs to their host wasp has long suggested to researchers that expansion of the virus gene families should result in altered host susceptibility (33, 34). Such susceptibility may take the form of additional susceptible hosts, increased numbers of pathways affected, or multiple tissues affected. This potential differential pathology has not been systematically analyzed functionally, in part due to the recalcitrance of PDVs to experimental manipulation and analysis: PDVs have multiple genes per gene family (35), with dynamic expression profiles that may differ across hosts (18, 19), and there currently is no *in vitro* system to streamline genetic manipulation for high-throughput study. Given the breadth of the potential impacts of genes and the difficulty of genetic screening, it is not surprising that few PDVs have been systematically analyzed.

Genome duplication in viruses is predicted to result in loss or novel functions where it provides for selective advantage (36). Previous studies with PDVs have suggested functional novelty or sub-functionalization: there are transcriptional differences seen between pathogenic hosts for a particular PDV, such as with HdIV (18), DfIV (19) and CsIV (34), and protein expression differences between host tissues, such as CcIV vankyrin (37) and MdBV PTPases (38), implying evolutionarily significant sub-functionalization. However, it is important to empirically test for potential sub-functionalization in an appropriate physiological context to determine potential contribution of the individual gene family members to host suitability. To that end, we tested two CsIV Vinnexins for their pathogenic contribution in a host, *H. virescens*, particularly in light of previously characterized pathophysiology noted in heterologous systems (27, 28).

We utilized recombinant baculoviruses to drive transgene expression in caterpillar hosts. We observed cellular localization (Fig. 1) reminiscent of VnxQ2 in CsIV-infected caterpillars (24), as well as VnxG and VnxQ2 expression in cell culture (27, 28). We observed increased mortality (Fig. 2) and reduced molting (Fig. 3) of the *vinnexin*- recombinant-infected larvae over the mock and recombinant virus controls. Previous studies with injections of purified CsIV have described increased mortality and molt-inhibition (39). Developmental inhibition has been linked to alterations in Juvenile Hormone/Ecdysone titers (40), which in turn have been associated with reduced prothoracicotropic gland size and activity (41). CsIV vankyrin expression in *Drosophila melanogaster* using the GAL4/UAS system indicates that single members of that gene family are sufficient to induce prothoracic gland cell dysfunction (42). We have not tested *vinnexin* expression or protein effect on endocrine gland function or circulating hormone titers. However, while the Vinnexins have inconsistent effects on cell number [Fig. 4; (27, 28)], it seems more likely that the molt pathology associated with them is general rather than targeted for the developmental system, as transcription of all four *vinnexins* is detected in all tested tissues in infected *H. virescens* (24). To that end, we suspect that the developmental delay observed here is due to cryptic cell death and loss of homeostasis, rather than pathology specific to the molting axis.

We also recapitulated previous *in vitro* observations of Vinnexin-specific alteration of membrane potential. Previously, we demonstrated that CsIV VnxG and VnxQ2 induce membrane depolarization and cytoplasmic alkalization in expressing Sf9 cells (27). Here, we found that hemocytes isolated from individuals infected with the vnxG-recombinant virus were depolarized relative to those isolated from vnxQ2-recombinant infected individuals, which in turn were depolarized relative to mock and control recombinant-infected hemocytes (Fig. 5). Intriguingly, little examination of the relationship between hemocyte function and membrane potential, as well as with endogenous and exogenous electrical fields, has been performed in insects. Caterpillar hemocytes undergo depolarization during immune stimulation (43), although the mechanism(s) and consequences of this are unknown. Additionally, disruption of the SERCA pump by venom components in hemocytes of *Leptoplina*-parasitized *D. melanogaster* larvae inhibits host immunity (44). While circumstances suggest that the Vinnexin-induced depolarization is important to hemocyte pathology, the mechanisms underlying it are currently unknown (27), and both mechanism and implications are under study.

It is now obvious that the Vinnexins have sub-functionalized: the electrophysiological properties associated with Vinnexin gap junctional intercellular communication, including apparent ability to form functional channels with the host Inx2 protein, differs between Vinnexins (24, 26), and the behaviors of cells expressing the vinnexins change in a protein-specific fashion both *in vitro* (27, 28) and *in vivo* (here). Importantly, there are functional consequences to these differences that suggest multimember families such as the vinnexins proliferate under pressure to complement one another, rather than to provide redundancy. VnxG and VnxQ2 both alter immune responses in expressing caterpillars relative to mock and recombinant virus controls (Fig. 6). Intriguingly, while VnxG disrupts encapsulation resulting in under- or over-formation of capsules, hemocytes from vnxQ2-infected individuals typically failed to form a capsule. It is possible that the two proteins differ due to inability of one to interact with *H. virescens* cellular machinery. While multiple virus hosts were not tested here, previous work has demonstrated consistency in pathological effect across numerous heterologous systems: VnxG forms stronger, more reliable gap junctions in *Xenopus* oocytes (24, 26), its expression is embryonic lethal in *D. melanogaster* embryos while VnxQ2 is not (28) and VnxG is more strongly alkalizing and depolarizing in the *Spodoptera frugiperda*-derived Sf9 cell line (27). Reduced pathogenicity is observed with VnxQ2 in these different systems, suggesting that while VnxQ2 is competent for interacting with host cells in a consistent fashion, it does so to a lesser effect than VnxG. While the question remains as to the evolvability of the system, it does suggest that VnxQ2 is less pathogenic than VnxG has evolved to be.

We performed descriptive and functional assays representing multiple physiological systems within the host *H. virescens* for two members of the CsIV *vinnexin* gene family. We observed significant variation in functionality between VnxG and VnxQ2, which is consistent with direction when compared to other tested systems. Our results thus suggest that the Vinnexins interact consistently and reliably between systems across hosts. However, it remains to be tested whether Vinnexins equally affect different hosts, which will be essential to address their potential role in evolution of host range and physiological susceptibility.

## 6. Methods

### Insect Rearing

Third instar *H. virescens* caterpillars were purchased (Benzon Research, Carlisle, PA) and reared at 27°C. Newly molted 4th instar larvae, staged according to head capsule width (45), were used for experiments, and were maintained at 27°C after treatment for data collection.

### Recombinant Virus Generation and Injection

The recombinant baculoviruses used for *in vivo* expression of *vnxG*-His and *vnxQ2*-His or control virus, FLAG-*mc4r*, were generated with Bac-to-Bac vector system (Invitrogen). Details for generation were previously published (27). Viruses were titered by plaque or end-point dilution assay (46). For injections, caterpillars were anaesthetized by submerging in ddH_2_O for 20 min and then the abdominal skin was sterilized with 70% ethanol and dried with paper towel. Five microliters Hink’s TNM-FH Insect Medium containing no recombinant virus or 100 pfu recombinant virus (*mc4r*, *vnxG*, *vnxQ2*) were injected laterally by a Hamilton #701 needle. A thin layer of liquid bandage (New-skin, Moberg Pharma North America LLC) subsequently was applied to the injection site to stop bleeding.

### Immunoassays

To collect hemocytes, caterpillars (5 caterpillars per treatment) at 3 days post-treatment were immobilized on ice and bled directly into cold anti-coagulant buffer (0.098M NaOH, 0.186M NaCl, 0.017M EDTA, 0.041M Citric acid, pH 4.5). Hemolymph samples were immediately centrifuged for 5 min at 500 × g, 4 °C. The supernatant was discarded and the hemocyte pellets were gently rinsed twice with cold PBS (pH 7.0). For western blot, hemocytes were resuspended in lysis buffer (25mM Tris-HCl, pH 7.6; 150Mm NaCl; 1% NP-40; 0.5% TritonX-100; 0.1% SDS). Equal concentrations of total protein, as determined by Bradford assay, were diluted in 4X loading buffer, incubated for 30 min at 37 °C, separated on 10% polyacrylamide gels (Bio-Rad, Mini-PROTEAN_®_ TGX™ Precast Gels) and transferred to PVDF membrane. Blots were probed with rabbit anti-His antibody (Thermo Fisher Scientific) at 1: 1,000 in blocking solution at 4 °C overnight, and then probed with polyclonal donkey anti-rabbit HRP-antibody (Invitrogen) at 1: 5,000 in blocking solution at room temperature for 1h. After washing, blots were treated with ECL substrate (Thermo Fisher Scientific), developed and visualized on X-ray film.

For immunomicroscopy, 10_4_ infected hemocytes was seeded in chamber slide, fixed with 4% formaldehyde (in PBS) at room temperature for 15 min, permeabilized with PBST (PBS + 0.2% TritonX-100) at room temperature for 10 min and blocked with 5% FBS (in PBST) at room temperature for 60 min with washing in PBS between every step. Primary (rabbit anti-His, Thermo Fisher Scientific) and secondary (anti-rabbit Alexa Fluor 594, Jackson ImmunoResearch) antibodies were diluted in blocking solution and applied at 1: 200 and 1: 1,000, incubated at 4 °C overnight and room temperature for 1 h, respectively. Images were captured on a Nikon TE2000 epifluorescence microscope with NIS Elements BR 2.3 software.

### Caterpillar Survival and Development

Thirty newly molted 4th instar larvae were injected with media or recombinant virus as above, and monitored daily for mortality for one week. Caterpillars were recorded as live, dead or pupated. Dead caterpillars were removed from diet tray after counting to avoid contamination. Larvae that died during the first three days were considered as mortality due to physical damage, and were removed from analyses. The instar of each caterpillar was recorded at 3 dpi and the proportion of 4th instar caterpillar was calculated. Each treatment was replicated 3 times.

### Hemocyte Count Determination and V_mem_ Measurement

Hemocytes were collected and washed as described above. Hemocyte counts were determined with a Neubauer hemocytometer, with three replicates per treatment. To measure the membrane potential of hemocytes, 10_4_ hemocytes were seeded in a 96-well plate and given 15 min to attach. At that time, Bis-(1,3-Dibutylbarbituric Acid) Trimethine Oxonol (DiBAC4(3)) was added to 1 μg/ml and allowed to incubate for 10 min in the dark. Hemocytes were imaged and analyzed as previously described (27). In brief, images were captured with a DS-Qi1Mc monochromatic camera on a Nikon TE2000 epifluorescence microscope with FITC filters. Cell intensity was collected by randomly choosing individual cells and manually generating the outline of cells as a Region of Interest (ROI). Normalized mean intensity (NMI) was calculated for each ROI by normalizing the ROI mean intensity (RMI) to background (BACK) and ambient light (AMB) by the calculation, NMI = [(RMI-BACK) / (AMB-BACK)] (27). A minimum of 165 cells was analyzed across three replicates for each treatment.

### *Ex Vivo* Encapsulation Assay

To test ability of hemocytes to engage in encapsulation, DEAE Sephadex beads were used as targets in ex vivo encapsulation assay. One million hemocytes of from treatment were placed in a 48-well plate coated with a layer 0.5% sterilized agarose (47). Ten beads, hydrated and sterilized, were added to each well. After 3 h incubation, beads were collected by gentle pipetting and transferred to a slide and only intact beads were examined for encapsulation. Beads were scored as “Normal encapsulation” if they were completely surrounded by multiple layers of hemocytes. “Abnormal encapsulation” noted both partial encapsulation- 75% surface was surrounded by hemocytes and as intense encapsulation- the encapsulation was several times as big as bead itself. Immune assay was repeated 5 times and at least 21 beads were examined for each treatment.

### Statistical Analyses and Figures

All statistical analyses were performed in R ×64 3.4.3 and Minitab 18. Graphs were generated in DataGraph V4.2.1 (Visual Data Tools Inc.).

## Abbreviations

PDV: Polydnavirus
IV: Ichnovirus
CsIV: *Campoletis sonorensis* Ichnovirus
inx: *innexin*
vnx: virus *innexin*
V_mem_: membrane potential

## Acknowledgements

This work was supported in part by USDA-AFRI award #2006-03787 and Clemson Creative Inquiry funding (Project #810). PZ and MT designed experiments, which PZ carried out. The authors would like to thank Richard Melton, Daniel Howard and Jessie Parker for assistance in managing larvae and for feedback.

